# A scalable method for preparation and long-term preservation of tea leaf plant-derived vesicles in tissue regeneration and drug encapsulation

**DOI:** 10.64898/2026.11.15.766803

**Authors:** Miriam M. Rodríguez de Lope, Alan Rivera-Tenorio, Ángel Moreno-Luque, Eduardo Chicano-Gálvez, Fernando de la Cuesta

**Affiliations:** Department of Pharmacology, School of Medicine, Universidad Autónoma de Madrid, Spain; Instituto de Investigación Hospital Universitario La Paz (IdiPaz), Madrid, Spain; Instituto de Medicina Regenerativa Tisular (IMERETI S.L), Madrid, Spain; IMIBIC Mass Spectrometry and Molecular Imaging Unit, Maimonides Biomedical Research Institute of Cordoba (IMIBIC), Reina Sofia University Hospital, University of Cordoba (UCO), Córdoba, Spain

## Abstract

Plant-derived vesicles (PDVs) are emerging as sustainable nanotherapeutic platforms, although challenges regarding isolation, scalability, biological activity, drug loading, and storage stability hinder their biomedical translation. This study aimed to optimize PDV production, exploring different sources to assess the one with greater translation potential, in terms of for yield and purity. To that end, PDVs from tea and red cabbage leaves, as well as ginger and turmeric roots were isolated by differential ultracentrifugation and size-exclusion chromatography (SEC), comparing both methodologies. Tea provided the highest vesicle yield and purity, with SEC outperforming ultracentrifugation in reducing soluble protein contamination. SEC isolated tea leaf-derived PDVs (Tea PDVs) were therefore further optimization using scalable fast protein liquid chromatography. Proteomic characterization showed enrichment in hydrolases, which could be related to an inherent capacity to induce tissue remodelling. Hence, regenerative potential was tested *in vitro* on mesenchymal stem cells (MSCs) and endothelial cells (HAECs). Firstly, Tea PDVs showed no cytotoxicity in MSCs and HAECs, even at high concentrations. Importantly, they significantly attenuated staurosporine-induced MSC death and promoted endothelial wound closure and capillary-like network formation, supporting their regenerative potential. Drug delivery potential was also assayed using quercetin, as a proof-of-concept drug. Saponin-mediated permeabilization significantly improved quercetin loading compared with passive incubation. Another important hurdle towards clinical translation, is determination of the ideal storage conditions for PDVs, which is still missing. To address this, we performed a six-months storage study at −80 °C, analysing different cryopreservatives. The results showed that PBS alone caused substantial particle loss and increased vesicle size, whereas HEPES + trehalose-, sucrose-, and trehalose + human albumin-containing cryoprotective formulations preserved vesicle size and mitigated particle loss, facilitating long-term stability.

Overall, these findings identify Tea PDVs as promising biocompatible nanocarriers with intrinsic regenerative properties. The optimized isolation, drug-loading, and preservation strategies provide a foundation for further development of plant-based nanotherapeutics targeting tissue regeneration and drug delivery applications in regenerative medicine.

## INTRODUCTION

Over the past decade, the field of nanomedicine has undergone a paradigm shift with the emergence of plant-derived vesicles (PDVs) as a promising and sustainable alternative to both mammalian extracellular vesicles and synthetic nanocarriers, such as liposomes. Although these vesicles have been described using different terms, including exosome-like nanovesicles, plant-derived nanovesicles or plant extracellular vesicles, recent efforts led by the International Society for Extracellular Vesicles (ISEV) Plant EVs Task Force aim to harmonize their nomenclature, providing scientific rigor in acknowledging the origin and purity of the vesicle isolates obtained. Accordingly, the term plant-derived vesicles (PDVs) will be used throughout this manuscript. These natural nanostructures, characterized by a lipid bilayer resembling that of mammalian EVs (Alzahrani et al., 2023), offer several advantages, including scalable and cost-effective production, reduced ethical concerns, excellent biocompatibility, and a favorable safety profile owing to their origin from edible plants (Logozzi et al., 2022; Wang et al., 2024; Zhuang et al., 2016). Similar to mammalian EVs, PDVs can serve as drug delivery vehicles while also carrying intrinsic bioactive cargo, including lipids, proteins, and microRNAs, with therapeutic effects (Alzahrani et al., 2023; Nemati et al., 2022). Indeed, vesicles isolated from edible plants such as ginger, grapefruit, lemon, and many others have demonstrated potent anti-inflammatory, antioxidant or antitumor activities by modulating key biological pathways(Chen et al., 2019; Raimondo et al., 2015; Zhuang et al., 2016). Consequently, PDVs can be now recognized as “green” nanotherapeutic platforms that combine the tradition of herbal medicine with modern biotechnology.

The biological properties of PDVs are largely determined by the molecular composition inherited from their plant of origin (Nemati et al., 2022). Consequently, the selection of the source plant is an important factor that can influence vesicle yield, molecular composition and biological activity.

Despite the growing number of studies describing PDVs from different edible plants, there are no direct comparative analyses between different plant sources regarding vesicle isolation yield, which represents one of the first challenges in the development of this field. Moreover, although the scalability of PDVs is frequently highlighted as one of their main advantages over mammalian EVs (Logozzi et al., 2022), preservation of vesicle integrity and functionality during long-term storage needs to be guaranteed for efficient therapeutic application. However, there are few studies evaluating the stability of PDVs and there is a lack of standardized strategies to improve their preservation. Likewise, while numerous reports have explored the use of PDVs as drug delivery vehicles, relatively few have optimized drug-loading procedures or quantitatively assessed loading efficiency, making it difficult to maximize their therapeutic potential.

Beyond these technological challenges, a better understanding of the biological interactions of PDVs is also required to support their biomedical translation. To date, most studies have focused on tumor models or inflammatory disease models (Cao et al., 2025; Chen et al., 2019; Raimondo et al., 2015; Zhuang et al., 2016), whereas their interaction with healthy cells involved in tissue repair and regeneration has received little attention. In particular, their effects on endothelial and mesenchymal stromal/stem cells, two key cellular types that play essential roles in angiogenesis and tissue regeneration, remain unexplored.
In this study, we selected four edible plants with well-documented therapeutic potential but distinct phytochemical profiles: tea (*Camellia sinensis*), red cabbage (*Brassica oleracea var. capitata f. rubra*), ginger (*Zingiber officinale*), and turmeric (*Curcuma longa*). These plants are rich in different classes of bioactive compounds, including catechins, anthocyanins, gingerols, and curcuminoids, respectively (Cao et al., 2025; Chen et al., 2019; Gao et al., 2022; Zu et al., 2021). To identify the most suitable source for the remainder of the study, PDVs were initially isolated from all four plant species using ultracentrifugation (UC) and size exclusion chromatography (SEC), allowing a comparative evaluation of their isolation yield and purity. Tea-derived PDVs (Tea PDVs) showed the highest yield and purity and were therefore selected for the remainder of the study. They were subsequently purified at a larger scale by bind-elute SEC on a fast protein liquid chromatography (FPLC) system, achieving a remarkable purity, as measured by PDV/protein ratio, and characterized in terms of proteomic composition, which pointed to a potential role in tissue regeneration. Based on these findings, and after first confirming their biocompatibility, we assessed their regenerative potential in endothelial cells and MSCs through functional assays of apoptosis, wound healing and angiogenesis. We also optimized a drug-loading strategy to improve encapsulation efficiency. Finally, since this work aims to evaluate Tea PDVs as a “green” platform for regenerative medicine applications, we assessed their long-term stability at ™80 °C and investigated stabilization strategies to preserve vesicle integrity and minimize particle loss, in order to maximize their therapeutic potential.

## MATERIALS AND METHODS

### Isolation of plant-derived vesicles (PDVs)

Fresh tea leaves (*Camellia sinensis*) were collected from plants maintained in our facilities, whereas fresh red cabbage (*Brassica oleracea var. capitata f. rubra*), ginger (*Zingiber officinale*) rhizomes, and turmeric (*Curcuma longa*) rhizomes were purchased from a local supermarket. All plant materials were surface-sterilized with 70% ethanol, rinsed twice with cold phosphate-buffered saline (PBS), and homogenized in cold PBS using a OV 625 Digital Dispersor homogenizer (Velp Scientific). The homogenate was sequentially filtered through a 70 μm cell strainer (Cultek) and centrifuged at 2,000 × g for 10 min at 4 °C, followed by 10,000 × g for 10 min at 4 °C to remove cell debris. The resulting supernatant was subsequently filtered through 0.45 μm and 0.22 μm syringe filters (Jet Biofil).

PDVs were isolated using either size exclusion chromatography (SEC), differential ultracentrifugation (dUC) or FPLC. For each *in vitro* experiment, the PDVs were freshly prepared. For SEC isolation, the clarified supernatant was concentrated 15–25-fold by ultrafiltration using Amicon Ultra-15 centrifugal filters (100 kDa molecular weight cut-off; Millipore, Merck) at 3,500 × g and 4 °C. Concentrated samples were loaded in in-house casted Sepharose CL-2B agarose 10 ml columns (Sigma-Aldrich), and 10 ml fractions were collected. Soluble protein contamination was evaluated using a bicinchoninic acid (BCA) protein assay, and fractions containing less than 0.05 μg/μL soluble protein were pooled and concentrated using Amicon Ultra-15 centrifugal filters (10 kDa molecular weight cut-off; Millipore, Merck). For dUC isolation, clarified supernatants were transferred into ultracentrifuge tubes (Beckman Coulter) and centrifuged at 100,000 × g for 1 h at 4 °C using an L8-70M ultracentrifuge (Beckman Coulter). Pellets were washed with PBS and a subsequent ultracentrifugation step was carried out under the same conditions, finally resuspending the pellet in PBS to the desired volume. The final preparations were passed through a 0.22 μm filter (Jet Biofil), and residual soluble protein contamination was assessed by BCA assay.

For bind-elute SEC, Tea PDVs were subsequently purified by fast protein liquid chromatography (FPLC) using an ÄKTA Start system equipped with a Capto Core 700 column. PDV-enriched fractions were either used directly or further concentrated, when necessary, using Amicon Ultra 100 kDa filters to the desired volume. Final preparations were stored at ™80 °C until characterization and functional analyses.

### PDVs characterization

Particle concentration, mean diameter, mode and size distribution of PDVs were determined by nanoparticle tracking analysis (NTA) using a NanoSight LM10 instrument (Malvern Panalytical, UK) equipped with a NanoSight Syringe Pump to ensure a constant sample flow during acquisition. Five technical measures from 60 seconds video captures were performed for each biological replicate.

Morphology of Tea PDVs was examined by transmission electron microscopy (TEM) using a JEM-1010 electron microscope (JEOL, Peabody, MA, USA). Samples were fixed with 2% paraformaldehyde for 30 min, deposited onto Formvar-carbon-coated copper grids (Ted Pella), negatively stained with 2% uranyl acetate and 0.5% methylcellulose.

### Proteomic analysis

Proteomic analysis of Tea PDVs was performed following the workflow previously described by our group (Rodríguez De Lope et al., 2025), with minor modifications. Briefly, PDVs samples were lysed, proteins were extracted and quantified, and tryptic peptides were generated using the iST kit (PreOmics, Germany). Peptides were analyzed in triplicate by liquid chromatography–tandem mass spectrometry (LC–MS/MS) using an Evosep One system coupled to a timsTOF Flex mass spectrometer (Bruker Daltonics) operating in DIA-PASEF mode. Protein identification and label-free quantification were performed using Spectronaut (Biognosys) with the *C. sinensis* UniProt database (June 2023). Functional enrichment analyses and protein-protein interaction networks were carried out using STRING database with default parameters, using ‘high confidence’ (0.700) as the minimum required interaction score, and hiding the unconnected nodes.

### Cell culture

Human telomerase reverse transcriptase (hTERT)-immortalized human adipose tissue-derived mesenchymal stem cells (AT-MSCs; ATCC) were cultured in DMEM (Corning) containing 4.5 g/L glucose and supplemented with 10% fetal bovine serum (FBS), 100 U/mL penicillin, 100 μg/mL streptomycin, and 2.5 μg/mL amphotericin B. hTERT-immortalized human aortic endothelial cells (TeloHAEC; ATCC) were cultured in M199 (Corning) supplemented with 25 µg/mL of endothelial cell growth supplement (ECGS; Sigma-Aldrich), 10% FBS, 100 U/mL penicillin, 100 μg/mL streptomycin, and 2.5 μg/mL amphotericin B. All cells were maintained at 37 °C in a humidified atmosphere containing 5% CO_2_. Unless otherwise indicated, culture medium was replaced every 48 h.

Cells were grown to the desired confluence and then cultured in medium containing extracellular vesicle-depleted FBS (prepared by ultracentrifugation at 100,000 × g).

### Citotoxity assays

The potential cytotoxicity of Tea PDVs was evaluated by crystal violet assay in AT-MSCs and TeloHAEC cells. Cells were seeded in 24-well plates at a density of 8,000 cells per well and incubated for 24 h to allow cell adhesion. Cells were subsequently starved overnight, with AT-MSCs maintained in medium supplemented with 1% extracellular vesicle-depleted FBS and TeloHAEC cells maintained in medium supplemented with 10% extracellular vesicle-depleted FBS without ECGS. Following the starvation period, cells were treated with Tea PDVs at doses of 50,000, 100,000, and 250,000 vesicles per cell. Each experimental condition was performed using three biological replicates, with each replicate analyzed in technical triplicate.

Following treatment, cell viability was determined by crystal violet staining. The culture medium was removed, and cells were washed with PBS. Cells were then fixed with 500 µL of 1% glutaraldehyde (VWR Chemicals) for 10 min at room temperature and washed three times with PBS. Subsequently, 500 µL of 1% crystal violet solution (Merck) was added to each well and incubated for 30 min at room temperature. Following staining, the wells were carefully washed with distilled water to remove excess dye and the plates were allowed to air-dry. The retained dye was solubilized by adding 2 mL of 10% acetic acid (Panreac) to each well and incubating for 5 min at room temperature. After homogenization, 250 µL from each well was transferred in technical duplicate to a 96-well plate, using 10% acetic acid as blank. Absorbance was measured at 595 nm using a FLUOstar OPTIMA spectrophotometer.

### Anti-apoptosis assays

AT-MSCs were seeded in 48-well plates at a density of 2 × 10^4^ cells per well. After 24 h starvation using 1% extracellular vesicle-depleted FBS, apoptosis was induced with 0.125 μM staurosporine (STS; Merck) for 16 h. Simultaneously, some plates were treated with Tea PDVs at doses of 5,000 and 50,000 particles per cell. At the end of the incubation period, cell viability was assessed using the crystal violet assay as described above.

### Angiogenesis assays

The pro-angiogenic potential of Tea PDVs was evaluated using a tube formation assay with TeloHAEC cells. The assay was performed using µ-Slide 15 Well 3D ibiTreat plates with polymer coverslip bottom (ibidi). Briefly, 10 µL of Matrigel (Corning) was added to each well and plates were subsequently incubated for 30 min at 37 °C to allow complete polymerization. After that, 15,000 TeloHAEC cells were seeded per well in DMEM supplemented with 5% exosome-free FBS. Cells were incubated under basal conditions or treated with Tea PDVs at doses of 1,000, 5,000, or 20,000 vesicles per cell. A positive control treated with VEGF (50 ng/mL; Sigma-Aldrich) was included. All experimental conditions were performed in biological triplicate.

Tube formation was monitored periodically, and images were acquired approximately 4-5 h after cell seeding using an EVOS microscope (Thermo Fisher Scientific). The angiogenic response was qualitatively evaluated by visual inspection of the tubular structures formed under each experimental condition.

### Wound healing assays

Pro-migratory potential effect of Tea-PDVVs was evaluated by wound healing (scratch) assay in TeloHAEC cells. Cells were seeded in 24-well plates at a density of 2.5 × 10^5^ cells per well. Before cell seeding, two reference lines were drawn on the underside of each well to facilitate image acquisition at the same position throughout the experiment. After 8 h starvation in medium supplemented with 10% extracellular vesicle-depleted FBS without ECGS, a single linear wound was generated in the center of each well by gently scraping the cell monolayer with a sterile pipette tip. The wells were subsequently washed once with starvation medium to remove detached cells and cellular debris. Treatments were then added to starvation medium. Images were acquired immediately after the scratch was made (0 h) and at different time points of treatment. Four images for each time point and treatment condition were acquired per well along the wound.

A positive control treated with VEGF (50 ng/mL; Sigma-Aldrich) was included. All experimental conditions were performed in biological triplicate.

Wound closure was quantified from the acquired images at 0 and 16 h using Fiji/ImageJ. The percentage of cell-free wound area was determined using the Wound healing size tool for Fiji. The wound area was measured at each time point, and the percentage of cell free area was calculated relative to the initial wound area at 0 h.

### Drug loading of Tea PDVs

Tea PDVs (3 × 10^10^ particles/mL) were incubated with 20 μM quercetin (Q) in 3% DMSO. Passive loading was performed by incubating the mixture for 1 h at 37 °C under agitation. Active loading was evaluated using sonication, saponin (0.2, 0.5, 2 or 5%), or a combination of both methods. For sonication, samples were pre-incubated for 30 min at room temperature, followed by two sonication cycles (30 s and 1 min) with cooling on ice between cycles, and a final incubation for 1 h at 37 °C. For saponin-mediated loading, different concentrations of *Quillaja saponaria* saponin (Sigma Aldrich) were added and samples were incubated for 1 h at 37 °C. For the combined protocol, sonication was performed before saponin addition.

Drug loading was quantified after vesicle lysis with 1% CHAPS. Samples were dried under vacuum, resuspended in methanol, centrifuged (13,000 × g, 5 min), and analysed by mass spectrometer (MRM) coupled to UHPLC-MS/MS. Encapsulated Q was quantified using a calibration curve prepared in the presence of vesicles to discard matrix effects. Drug loading was expressed as the percentage of the initial Q concentration.

### Stability assays of Tea PDVs

To evaluate the stability of Tea PDVs during long-term storage, vesicles isolated by SEC were aliquoted and stored at ™80 °C under different buffer conditions for up to 6 months. Four storage conditions, consisting on 4 different conservation buffers, were tested: (1) PBS, (2) PBS supplemented with 25 mM HEPES, 0.2% human serum albumin, and 25 mM trehalose (Görgens et al., 2022), (3) PBS supplemented with 25 mM HEPES and 25 mM trehalose, and (4) PBS supplemented with 25 mM HEPES and 25 mM sucrose.

Particle concentration and average size were analyzed by NTA at 1, 2, and 6 months post-storage to assess vesicle stability over time.

## RESULTS

### Comparative isolation of PDVs from different edible plant sources identifies tea as the most suitable source for high-yield and high-purity PDV isolation

To identify the most suitable plant source for PDV isolation, vesicles were isolated from tea, ginger, red cabbage and turmeric using both SEC and dUC. For SEC, particle concentration and soluble protein content were determined in each fraction by NTA and BCA assay, respectively, allowing the identification of vesicle-enriched fractions with minimal protein contamination (Fig. 1A–D). In tea, red cabbage and turmeric, particle-enriched fractions were clearly separated from the main soluble protein peak, which allowed us to pool fractions containing less than 0.05 μg/μL of soluble protein for subsequent concentration and PDV recovery. In contrast, ginger-PDVs co-eluted with soluble proteins throughout the chromatographic profile, preventing the isolation of high purity vesicles by SEC, so ginger was therefore excluded from the SEC comparison.

**Figure 1.**
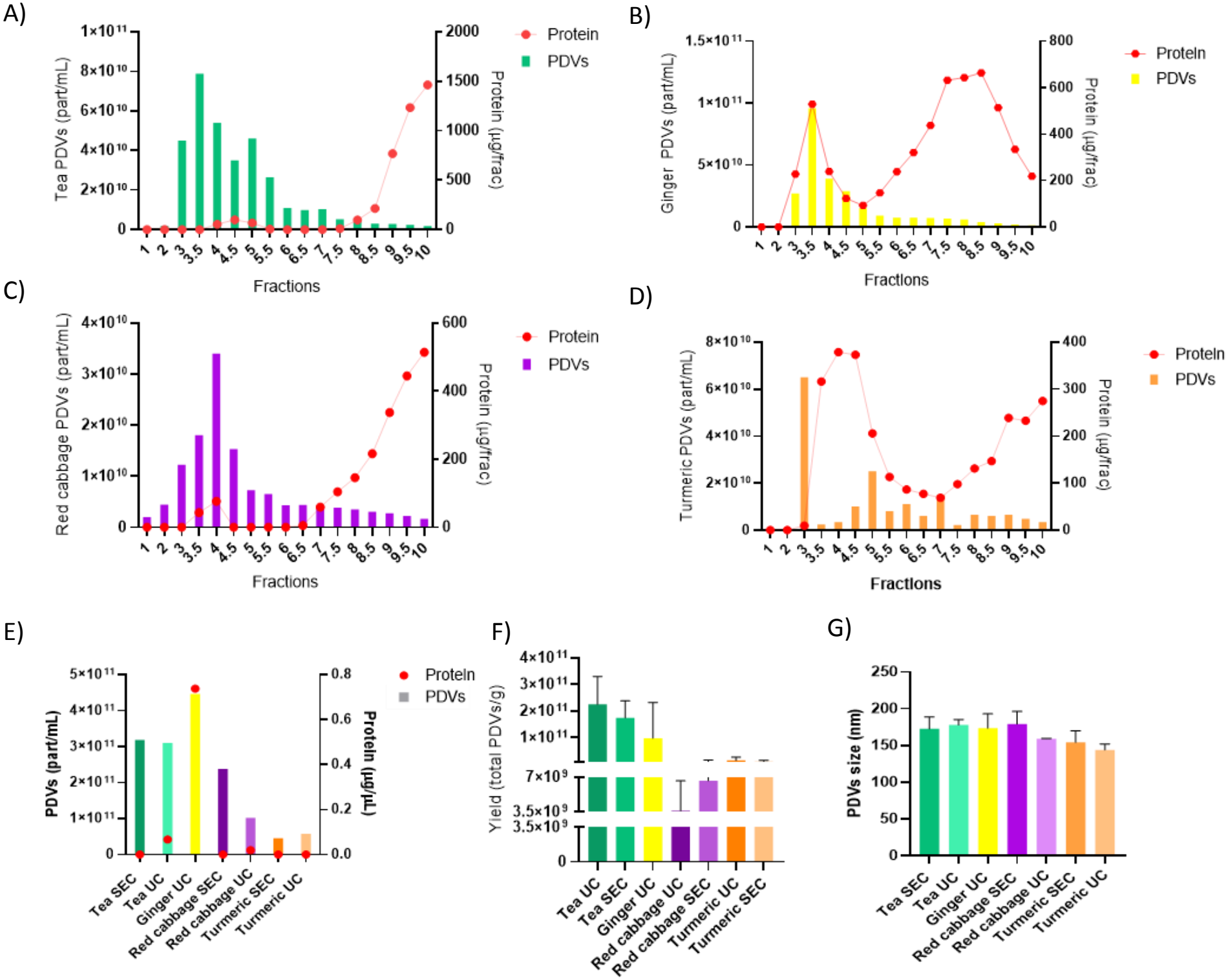
Comparative isolation of PDVs from tea, ginger, red cabbage and turmeric. (A–D) SEC elution profiles of PDVs from tea (A), ginger (B), red cabbage (C) and turmeric (D). Particle concentration (bars, left axis) was measured by nanoparticle tracking analysis (NTA) and soluble protein content (red dots, right axis) by bicinchoninic acid (BCA) assay in each fraction. (E) Particle concentration (bars, left axis) and soluble protein concentration (red dots, right axis) of the final PDV preparations obtained by SEC or dUC. Ginger-PDVs were isolated only by dUC, as SEC did not separate vesicles from soluble proteins. (F) Isolation yield, expressed as total PDVs per gram of starting material. (G) Mean PDV size determined by NTA. Data are presented as mean ± SEM of n =3.

Following concentration of the selected SEC fractions, PDVs preparations obtained by SEC and those obtained by UC were compared in terms of particle recovery, soluble protein contamination, yield and particle size (Fig. 1E–G). SEC preparations contained less soluble protein than dUC isolates, resulting in higher-purity PDV preparations [particles/μg protein] (Fig. 1E). Tea PDVs exhibited the highest particle yield with both isolation methods, whereas ginger-PDVs isolated by dUC yielded approximately half the number of recovered particles. In contrast, PDVs isolated from red cabbage and turmeric showed particle yields approximately one order of magnitude lower than those obtained from tea. Vesicle size remained comparable among all plant sources and was not affected by the isolation method (Fig. 1G). Overall, these results identified tea as the plant source with the best combination of particle yield and purity when PDVs were isolated by SEC, and Tea PDVs were therefore selected for all subsequent characterization and functional studies.

### Tea PDVs can be isolated at large scale by bind-elute SEC on an FPLC system

Following the selection of Tea PDVs isolated by SEC, the protocol was scaled up to a bind-elute SEC implemented on FPLC platform. This approach was evaluated as an alternative to conventional SEC, allowing a comparison of particle recovery, purity and processing time while providing a more standardized and automated isolation procedure suitable for large-scale production. The chromatographic fractions were characterized in terms of particle concentration, soluble protein content, and particle-to-protein ratio to assess the separation efficiency and identify the fractions with the highest purity (Fig. 2A). The analysis revealed differences in particle recovery and protein contamination across the collected fractions, allowing the selection of PDV-enriched fractions with reduced soluble protein content for subsequent pooling. We next evaluated the impact of each purification step on particle recovery and purity, from the initial crude plant extract through sequential filtration, tangential flow filtration (TFF), chromatographic purification, and final concentration by ultrafiltration (Fig. 2B). The progressive processing of the plant extract resulted in changes in particle concentration and soluble protein content, with chromatographic purification providing an improvement in the particle-to-protein ratio. The selected FPLC fractions were pooled and subsequently concentrated to obtain the final tea-PDV preparation. Nanoparticle tracking analysis (NTA) of the purified preparation revealed a predominantly nanosized particle population, with a modal diameter of approximately 140 nm (Fig. 2C). Transmission electron microscopy (TEM) further confirmed the presence of vesicular structures displaying rounded and cup-shaped morphologies, consistent with the characteristic appearance of extracellular vesicles (Fig. 2D).

**Figure 2.**
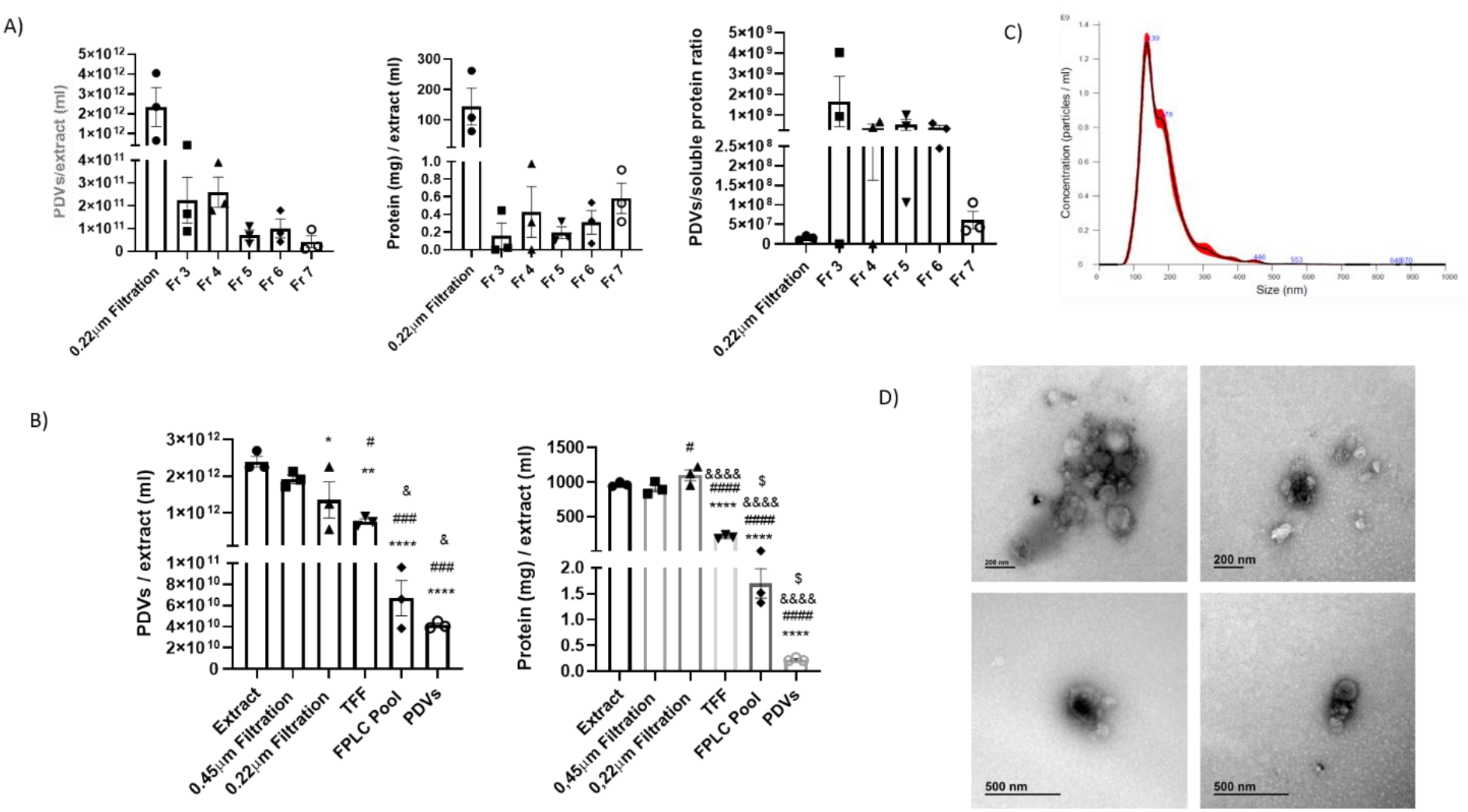
Optimization of large-scale isolation and purification of tea-derived vesicles (Tea PDVs) using bind-elute size-exclusion chromatography (BE-SEC) by fast protein liquid chromatography (FPLC). (A) Particle concentration (PDVs/mL), soluble protein concentration (mg/ml), and particle-to-protein ratio (PDVs/µg protein), determined in the initial sample loaded onto the FPLC system following 0.22 µm filtration and in the individual chromatographic fractions collected during BE-SEC purification. (B) Changes in particle concentration and soluble protein concentration throughout the sequential isolation and purification workflow, including crude extract preparation, two filtration steps (0,22 µm and 0,45 µm), tangential flow filtration (TFF), pooling of selected FPLC fractions (FPLC pool), and final concentration by ultrafiltration to obtain the purified tea-PDV preparation. (C) Representative nanoparticle tracking analysis (NTA) profile showing the particle size distribution and concentration of the final purified tea-PDV preparation. (D) Representative transmission electron microscopy (TEM) images of purified Tea PDVs, showing nanosized vesicular structures with rounded and cup-shaped morphologies. Scale bars: 200 nm (upper panels) and 500 nm (lower panels). Data are presented as mean ± SEM of n = 3 independent experiments. Statistical analysis was performed by one-way ANOVA followed by Dunnett’s multiple comparisons test.

### Proteomic analysis showing enrichment in hydrolases suggest potential to induce tissue remodelling

Tea PDVs were subjected to in-depth proteomic analysis, and 1623 proteins were identified. Functional enrichment analysis of the Molecular Function category (Fig. 3A) revealed a marked representation of proteins with enzymatic activity, predominantly hydrolases. This profile indicated that Tea PDVs contained a high proportion of proteins with catalytic functions associated with cellular degradation or remodelling processes, consistent with the potential contribution of components originating from the lytic vacuole, as found enriched in cell component category (Fig. 3B). Besides, the term secretory vesicle was also found significantly enriched (Fig. 3B). As many as 131 hydrolases were found in Tea PDVs, with several protein-protein interactions among them (Figure 3C).

**Figure 3.**
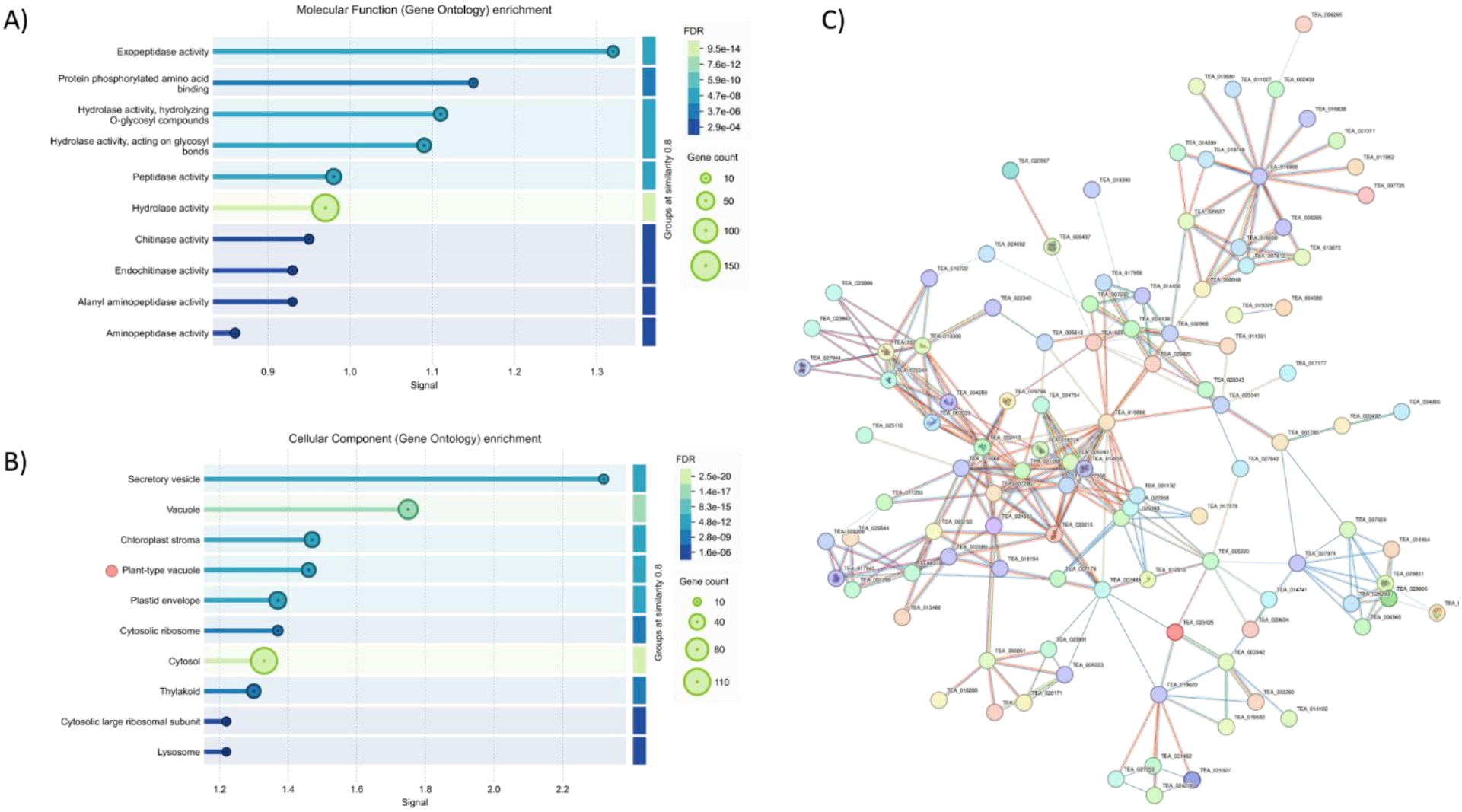
Proteomic characterization and functional enrichment analysis of tea-derived vesicles (Tea PDVs). A total of 1,623 proteins were identified by LC–MS/MS and subjected to functional enrichment and protein–protein interaction analyses using the STRING database. (A) Functional enrichment analysis of Gene Ontology (GO) Molecular Function terms, showing a predominance of proteins with catalytic activity, particularly hydrolases. (B) Functional enrichment analysis of GO Cellular Component terms, highlighting the significant representation of proteins associated with the lytic vacuole and secretory vesicles. In (A) and (B), bubble size represents the number of proteins associated with each GO term, while the color scale indicates the false discovery rate (FDR)-adjusted significance of the enrichment. (C) STRING protein–protein interaction network of the hydrolases identified in Tea PDVs which showed interactions (. Nodes represent individual proteins, while edges indicate known or predicted functional associations between proteins, with different colors corresponding to distinct sources of interaction evidence.

### Tea PDVs are non-cytotoxic, protect MSCs from apoptosis and promote capillary-like network formation on Matrigel and endothelial wound closure

After confirming that Tea PDVs can be isolated at large scale and stored with preserved size, we evaluated their biocompatibility and regenerative potential *in vitro*. To assess cytotoxicity, adipose tissue-derived MSCs (AT-MSCs) and telomerase-immortalized human aortic endothelial cells (TeloHAECs) were treated with increasing doses of Tea PDVs, up to 250,000 PDVs/cell, and cell viability was determined by crystal violet staining. Tea PDVs did not significantly affect the viability of either cell type at any dose tested (Fig. 4A).

**Figure 4.**
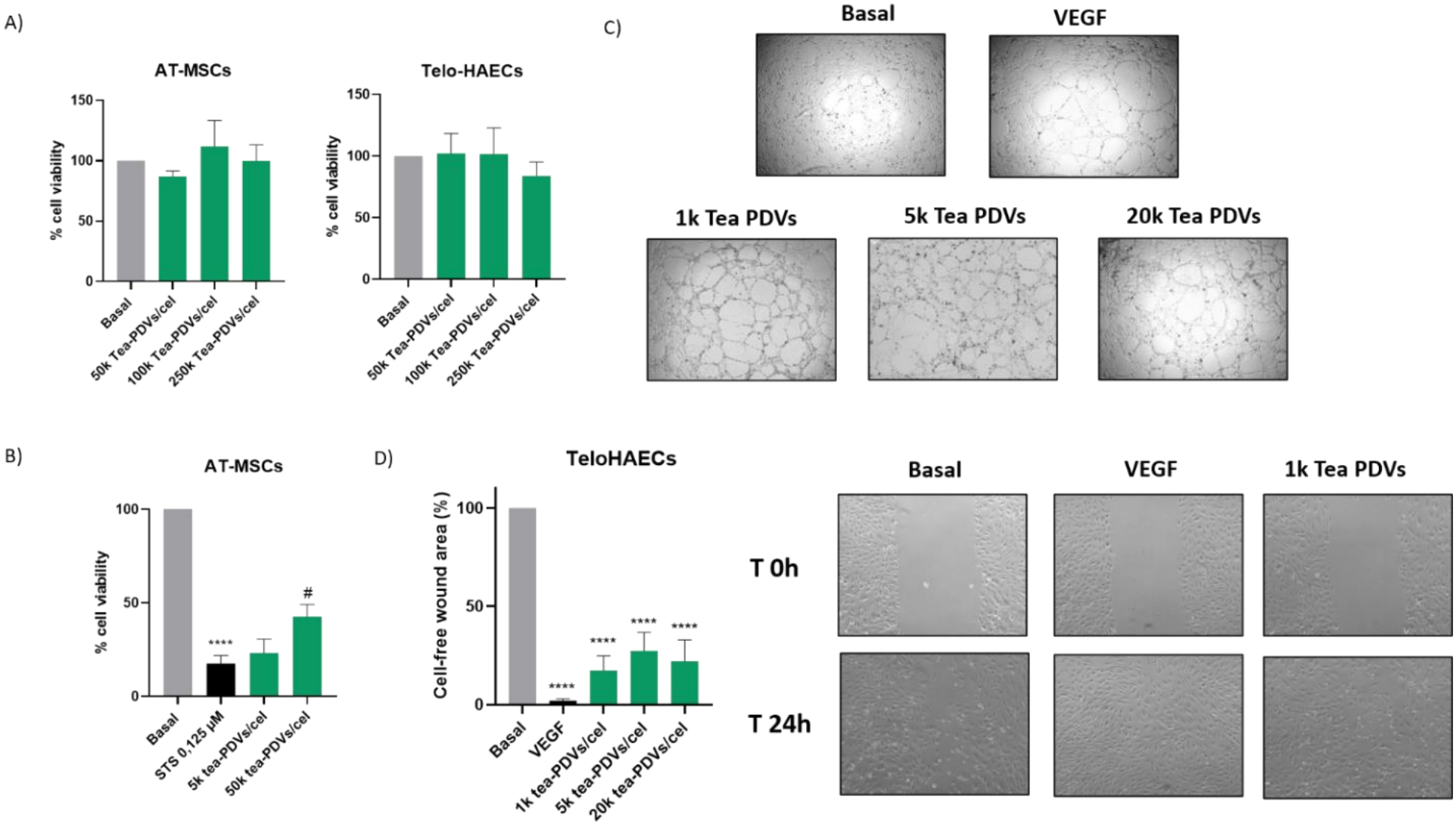
Biocompatibility and regenerative effects of Tea PDVs on AT-MSCs and TeloHAECs. (A) Viability of AT-MSCs and TeloHAECs treated with 50,000, 100,000 or 250,000 Tea PDVs/cell for 24h, determined by crystal violet staining and expressed as a percentage of basal (untreated cells). No significant differences were found versus basal. (B) Protection against staurosporine (STS)-induced apoptosis in AT-MSCs. Cells were co-treated with STS (0.125 μM) and 5,000 or 50,000 Tea PDVs/cell for 16h, and viability was determined by crystal violet staining and expressed as a percentage of basal. (C) Representative bright-field images of tube formation by TeloHAECs on Matrigel 4–5 h after seeding, in basal medium, with VEGF (50 ng/mL) or with 1,000, 5,000 or 20,000 Tea PDVs/cell. (D) Scratch wound healing assay in TeloHAECs under starvation conditions (without ECGS). Left, cell-free wound area after 24 h, expressed as a percentage relative to the initial wound area at 0 h. Right, representative images at 0 h and 24 h for basal, VEGF and 1,000 Tea PDVs/cell. Data are presented as mean ± SEM of n = 3 (A, B) and n = 5 (D) independent experiments. Statistical analysis was performed by one-way ANOVA followed by Dunnett’s multiple comparisons test (vs. basal in A, B and D; vs. STS in B). #P < 0.05 vs. STS; ^****^P < 0.0001 vs. basal.

We next explored whether Tea PDVs could protect cells from apoptosis, a property relevant for regenerative applications. For this purpose, AT-MSCs were treated with staurosporine (STS, 0.125 μM) alone or together with Tea PDVs. STS markedly reduced cell viability to 17.63 ± 4.13% relative to basal. Co-treatment with 50,000 Tea PDVs/cell partially attenuated this effect, increasing viability to 42.63 ± 6.51%, a significant increase compared with STS alone (*P < 0*.*05)* (Fig. 4B). The lower dose of 5,000 Tea PDVs/cell had no significant effect on viability, although a trend toward increased viability was also observed.To evaluate the effect of Tea PDVs on angiogenesis, we performed a tube formation assay on Matrigel using TeloHAECs. Cells treated with 1,000, 5,000 or 20,000 Tea PDVs/cell formed interconnected capillary-like networks 4-5 h after seeding, similar to those observed with VEGF (Fig. 4C).

Finally, the effect of Tea PDVs on endothelial cell migration was assessed by a scratch wound healing assay in TeloHAECs under starvation conditions. After 24 h, all doses of Tea PDVs significantly reduced the cell-free wound area compared with basal (1,000, 5,000 and 20,000 Tea PDVs/cell, all *P < 0*.*0001*; Fig. 4D). A dose of 1,000 Tea PDVs/cell was sufficient to markedly promote wound closure, reducing the cell-free area to 17.40 ± 7.43% of that of the basal condition. Higher doses (5,000 and 20,000 Tea PDVs/cell) did not further reduce the wound area (27.35 ± 9.44% and 22.08 ± 10.84% of basal, respectively), indicating no dose-dependent effect within the range tested. The positive control, VEGF, led to almost complete wound closure (2.04 ± 0.94% of basal). Overall, these results indicate that Tea PDVs are well tolerated by MSCs and endothelial cells, partially counteract apoptosis in MSCs and enhance endothelial wound closure.

### Saponin treatment improves quercetin loading into Tea PDVs

Finally, we evaluated strategies to load Tea PDVs with quercetin (QC). In previous work from our group, 0.2% saponin was the most effective active loading method for AT-MSC-derived EVs (unpublished, data not shown). Based on this, we tested higher saponin concentrations (0.2, 0.5, 2 and 5%), alone or combined with sonication, and compared them with passive incubation (1 h at 37 °C), with or without sonication, as controls. We assessed particle recovery, vesicle size and the amount of QC associated with the PDVs. These experiments were performed in two independent biological replicates and should therefore be considered exploratory.Particle recovery after passive incubation was low, both without and with sonication (32.42 ± 2.58% and 49.83 ± 14.83% of the initial particles, respectively), whereas saponin treatment markedly reduced particle loss, with or without sonication, with recoveries of 81-108% depending on the concentration (Fig. 5A). The exception was the combination of sonication with 0.2% saponin, which gave a lower and more variable recovery (52.23 ± 23.43%). Mean vesicle size remained close to baseline after treatment with saponin alone (94-104% of the initial size) and was moderately reduced when saponin was combined with sonication, down to 83.10 ± 6.1% with 5% of saponin (Fig. 5B).Regarding QC loading, passive incubation resulted in negligible amounts of QC associated with PDVs (<0.1% of the initial QC). Saponin alone increased QC loading, which tended to rise with saponin concentration, and was more effective than the combinations with sonication, particularly at 0.2% and 0.5% saponin (Fig. 5C). The highest loading was obtained with 5% saponin (4.96 ± 0.48% of the initial QC).

**Figure 5.**
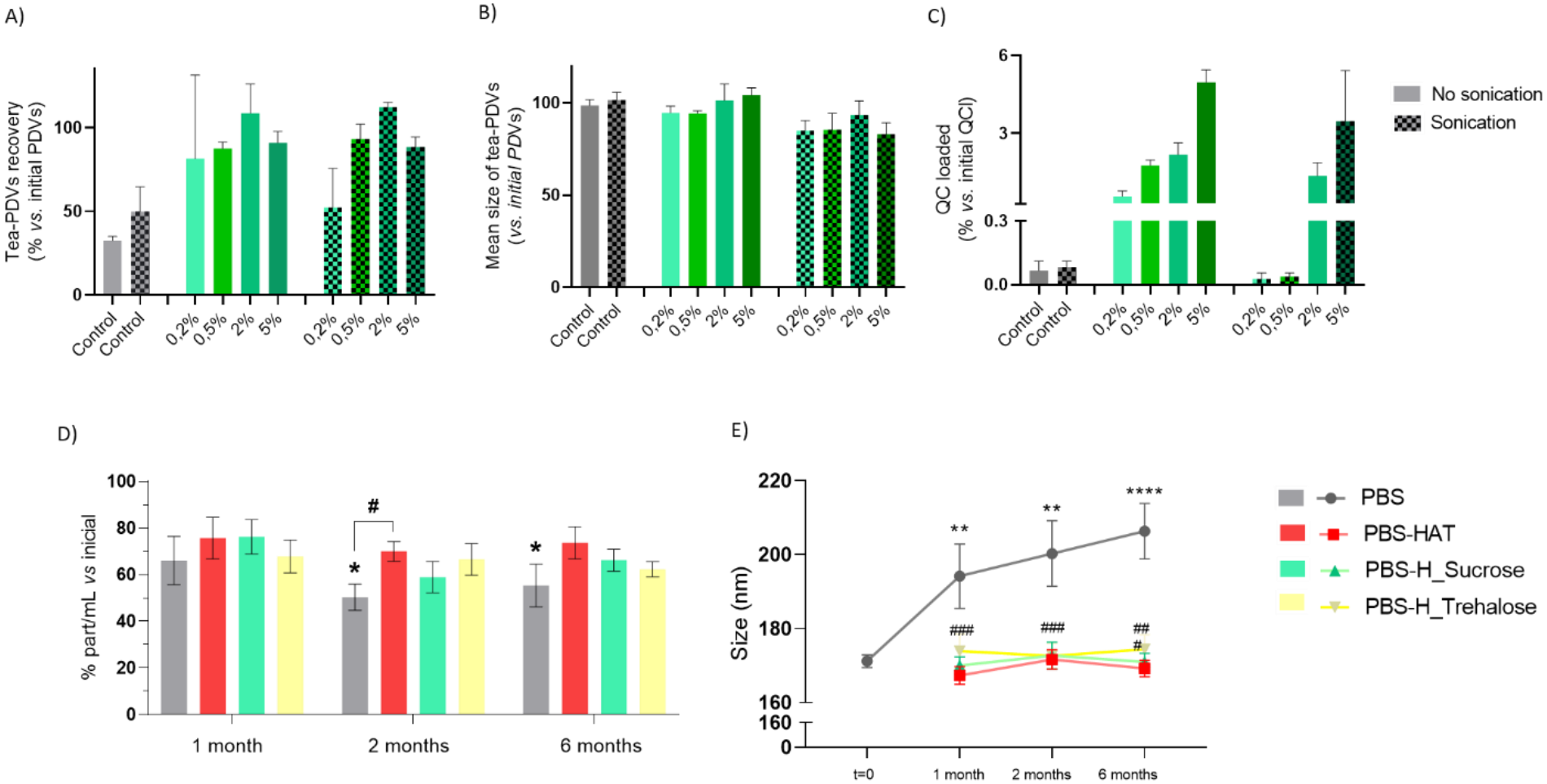
Quercetin loading and storage stability of Tea PDVs. (A–C) Tea PDVs were incubated with quercetin (QC) and 0.2-5% saponin, with or without sonication. Passive incubation (1 h, 37 °C) was used as control. Particle recovery (A) and mean size (B) were determined by NTA, and QC associated with PDVs (C) was quantified by UHPLC-MS/MS (MRM). All values are expressed relative to the initial values. Data are presented as mean of n = 2 ± SEM of independent biological replicates; no statistical analysis was performed. (D, E) Particle concentration, expressed as a percentage of the initial concentration (t = 0) (D), and mean size (E) of Tea PDVs measured by NTA after storage at -80 °C for 1, 2 or 6 months in PBS, Görgens buffer, 25 mM sucrose or 25 mM trehalose. Data are mean ± SEM of n = 3-6 independent isolations; unpaired Student’s t-test. In D, ^*^P < 0.05 vs. 1 month (same buffer); #P < 0.05 vs. PBS (same time point). In E, ^**^P < 0.01 and ^****^P < 0.0001 vs. t = 0; ###P < 0.001 vs. PBS (same time point).

### Cryoprotective formulations prevent the storage-induced size increase and partially attenuate particle loss of Tea PDVs

To improve the long-term preservation of Tea PDVs, vesicles were stored at ™80 °C in PBS or in the presence of three different cryoprotective formulations: trehalose (25 mM), sucrose (25 mM) or the Görgens preservation buffer previously described for mammalian EVs (Görgens et al., 2022), previously described for mammalian EVs storage, containing PBS, 25 mM HEPES, 0.2% human serum albumin (HSA) and 25 mM trehalose (Fig. 5D and E). In PBS, particle concentration decreased to 66,13 ± 10,31% of the initial value after one month and showed a further significant decrease to 50.41 ± 5.6% after two months, remaining at a similar level after six months (55.39 ± 9.16%; *P < 0*.*05* vs. one month) (Fig. 5D). In contrast, no significant decrease in particle concentration between one month and later time points was observed with any of the three cryoprotective formulations. Moreover, at two months, the Görgens buffer retained significantly more particles than PBS (Fig. 5D). Together, these results indicate that cryoprotective formulations partially mitigate the loss of PDVs during storage, although a reduction in particle concentration relative to the initial value was observed in all conditions.In addition to particle loss, storage in PBS induced a progressive increase in vesicle size, from an initial mean diameter of 171.25 ± 1.72 nm to 194.18 ± 8.66 nm after one month, 200.3 ± 8.88 nm after two months and 206.33 ± 7.49 nm after six months (Fig. 5E). This increase was completely prevented by all three preservation buffers, which maintained particle size close to baseline throughout the study.

## DISCUSSION

Lately, PDVs from edible plants are attracting increasing interest in biomedicine as a result of their availability, biocompatibility and low immunogenicity (Alzahrani et al., 2023), and the possibility of administering them through different routes (Wang et al., 2024). However, research on their isolation, characterization, biological function and capacity to carry biomolecules is still at an early stage. In this study, we compared four PDV sources from edible plant parts: tea and red cabbage leaves, and ginger and turmeric roots, identifying tea as the one providing the greatest yield of PDVs, and evaluated Tea PDVs as potential therapeutic agents and drug delivery vehicles, by assessing large-scale high-purity separation, proteomic composition, *in vitro* regenerative efficacy on MSCs and endothelial cells, drug loading and storage stability.

Among the studied plant sources, Tea PDVs showed the highest yield with both isolation methods, and SEC preparations contained less soluble protein than dUC isolates. Given that SEC is a separation method that better preserves vesicle integrity than dUC, the greater purity achieved made the selection of this procedure a clear decision. Of note, all other plant sources showed acceptable purity but ginger, in which our detailed analysis of soluble protein showed very poor purity in terms of PDV/protein ratio in both methods tested. In fact, selection of fractions with sufficient PDV purity was unsuccessful since none would follow our purity criteria.In a further step, the SEC method was scaled up using BE-SEC in an FPLC system, which allowed automating the process, while still providing a good yield and increased purity.

Before considering Tea PDVs for any further application, we confirmed their safety, as a prerequisite for their potential therapeutic use. Tea PDVs did not significantly affect the viability of AT-MSCs or TeloHAECs even at doses far above those used in the functional assays (250,000 PDVs/cell). Based on these proteomics results, we explored their regenerative potential using AT-MSCs, given their growing relevance in regenerative medicine (Keshtkar et al., 2018) and staurosporine-induced apoptosis as a cellular stress model (Kholodenko et al., 2022). Co-treatment with Tea PDVs partially attenuated the loss of viability induced by staurosporine with the highest dose. We also investigated their effects on endothelial cells, a well-established *in vitro* model for studying endothelial function (Goodwin, 2007; Nareshkumar et al., 2018). In the scratch assay, all tested doses of Tea PDVs reduced the cell-free area, suggesting a potential effect on wound closure. However, the absence of a clear dose-response relationship suggests that the effect saturates at low doses. In addition, tea-PDV-treated cells formed capillary-like networks in the tube formation assay, comparable to those observed following VEGF treatment. To our knowledge, this is the first time that a pro-angiogenic effect of Tea PDVs is described.

After characterizing these biological effects, we tested whether Tea PDVs could also be used as drug carrier. As a proof-of-concept, we selected quercetin, an anti-inflammatory and antioxidant flavonoid found in plants s (Deepika & Maurya, 2022). Passive loading under conditions previously optimized for AT-MSC-EVs gave markedly lower loading in Tea PDVs, which may reflect differences in membrane lipid composition and permeability (Alzahrani et al., 2023). Saponin alone, especially at 5%, gave the highest loading, about 70-fold above passive incubation, consistent with its permeabilizing action on animal EVs (Haney et al., 2015). Sonication did not improve loading and, combined with saponin, reduced vesicle size, probably through mechanical damage. Saponins are also considered biodegradable and may reduce the toxicity of free drugs (Liao et al., 2021). Further studies are needed to determine whether the amount of quercetin loaded into Tea PDVs is sufficient to produce relevant biological effects.

Finally, we decided to analyse one important aspect that has been overlooked in the field, the stability of plant vesicles overtime and the requirement for cryopreservatives. With this goal in mind, we examined their storage at -80 °C for six months in PBS alone, supplemented with a sugar: trehalose or sucrose, and the complete formulation described by Görgens et al. for MSC-EVs (HEPES, trehalose and human serum albumin). In PBS, recovery of Tea PDVs dropped up to around half of the initial number of particles, mostly during the first two months, additionally showing a substantial and progressive size increase, which suggests aggregation. The trehalose, sucrose and Görgens formulations prevented the size increase and showed no significant decrease in particle concentration after one month, consistent with their ability to prevent aggregation and freezing damage in EVs (Bosch et al., 2016; Charoenviriyakul et al., 2018; Janouskova et al., 2022). Differences in particle concentration between cryopreserving buffers were limited, with only the Görgens buffer retaining significantly more particles than PBS (at two months). The Görgens buffer contains human serum albumin, a clinically used excipient (Sattler et al., 2023), although its inclusion means that the final formulation will no longer remain entirely plant-derived. Trehalose and sucrose may therefore represent more suitable alternatives when maintaining a plant-compatible formulation is a priority, as they provided similar protection against changes in particle size and reduction in particle concentration was not statistically significant, as opposed to PBS alone. Although future studies should determine whether these formulations additionally preserve the biological activity of Tea PDVs during long-term storage, the preservation of PDV integrity and particle number are promising results in the application of these cryopreserving agents in the plant vesicle field.

In conclusion, this study identifies tea leaves as a promising source of high-yield, high-purity PDVs and establishes a scalable FPLC-based purification strategy. Proteomic characterization revealed an enrichment in hydrolases, suggesting potential involvement in tissue remodelling. Importantly, Tea PDVs demonstrated excellent biocompatibility, protected MSCs against apoptosis, and promoted endothelial wound closure, supporting their regenerative potential. Furthermore, saponin-mediated permeabilization enhanced quercetin loading, while cryoprotective formulations improved vesicle preservation during long-term storage. Together, these findings provide a foundation for the development of Tea PDVs as sustainable nanotherapeutic platforms combining intrinsic regenerative activity with drug delivery potential.

## Acknowledgements

The authors would like to thank Estela Herranz for her valuable technical assistance and support throughout this study, and Prof. Isabel Sánchez Pérez for her assistance with the wound-healing assay optimization.

